# Trophic cascade driven by behavioural fine-tuning as naïve prey rapidly adjust to a novel predator

**DOI:** 10.1101/856997

**Authors:** Chris J. Jolly, Adam S. Smart, John Moreen, Jonathan K. Webb, Graeme R. Gillespie, Ben L. Phillips

**Affiliations:** School of BioSciences, University of Melbourne, Parkville Vic 3010 Australia; Kenbi Rangers, Mandorah NT 0822 Australia; School of Life Sciences, University of Technology Sydney, Broadway NSW 2007 Australia; Flora and Fauna Division, Department of Environment and Natural Resources, Northern Territory Government, Berrimah NT 0828 Australia

**Keywords:** Antipredator behaviour, boldness, invasion, neophobia, novel predator, predator-prey dynamics, prey naivety

## Abstract

The arrival of novel predators can trigger trophic cascades driven by shifts in prey numbers. Predators also elicit behavioural change in prey populations, via phenotypic plasticity and/or rapid evolution, and such changes may also contribute to trophic cascades. Here we document rapid demographic and behavioural changes in populations of a prey species (grassland melomys *Melomys burtoni*, a granivorous rodent) following the introduction of a novel marsupial predator (northern quoll *Dasyurus hallucatus*). Within months of quolls appearing, populations of melomys exhibited reduced survival and population declines relative to control populations. Quoll-invaded populations (*n* = 4) were also significantly shyer than nearby, quoll-free populations (*n* = 3) of conspecifics. This rapid but generalised response to a novel threat was replaced over the following two years with more threat-specific antipredator behaviours (i.e. predator-scent aversion). Predator-exposed populations, however, remained more neophobic than predator-free populations throughout the study. These behavioural responses manifested rapidly in avoidance of seeds associated with quoll scent, with discrimination playing out over a spatial scale of tens of metres. Presumably the significant and novel predation pressure induced by quolls drove melomys populations to fine-tune behavioural responses to be more predator-specific through time. These behavioural shifts could reflect individual plasticity (phenotypic flexibility) in behaviour or may be adaptive shifts from natural selection imposed by quoll predation. Our study provides a rare insight into the rapid ecological and behavioural shifts enacted by prey to mitigate the impacts of a novel predator and shows that trophic cascades can be strongly influenced by behavioural changed rates of seed predation by melomys across treatments. Quoll-invaded melomys populations exhibited lower per-capita seed take rates, and rapidly developed an as well as numerical responses.

## Introduction

Predation is one of the most pervasive and powerful forces acting on populations. Not only does predation directly impact a population’s demography (Schoener & Spiller 1996), it also imposes natural selection (Abrams 2000). The pressure that predators impose on populations will vary through time and space for many reasons, including tightly coupled predator-prey dynamics, predator movement, prey switching, or stochastic processes (Lima & Dill 1990; Sih 1992). The fact that predation is not constant, and that antipredator defences may be costly, suggests that flexible responses to predation pressure will often be favoured (Sih *et al.* 2000; Berger *et al.* 2001). There is, in fact, a great deal of empirical evidence that flexible responses to predation are common (e.g. Relyea 2003; Brown *et al.* 2013; Cunningham *et al.* 2019). Investment in antipredator traits across morphology, life-history, and behaviour often varies, and is dependent on the perceived risk of predation.

As well as impacting prey populations, it is increasingly apparent that predators play a powerful role in structuring communities (Estes *et al.* 2011). Some of our best evidence for this comes from the introduction of predators to naïve communities. Invasive predators can cause extinctions (Medina *et al.* 2011; Woinarski *et al.* 2015; Doherty *et al.* 2016), and alter trophic structures and ecosystem function within recipient communities (Courchamp *et al.* 2003; Simberloff *et al.* 2013). Cascading outcomes are often thought of as purely numeric effects: predators depress the size of prey populations, and the altered numbers of prey can cause cascading numerical changes down trophic levels (Ripple *et al.* 2001). These numerical effects are undeniably important, but the fact that predators can also elicit phenotypic change in prey populations—through phenotypic plasticity and natural selection—means that subtler ecological effects may also manifest. Prey species living alongside predators may forage at different times, or in different places compared with the same species in a predator-free environment (Laundre *et al.* 2010). Such behavioural shifts can alter downstream species interactions in potentially complex ways (Fortin *et al.* 2005; Suraci *et al.* 2016).

Because predator invasions are rarely intentional or anticipated, there is a scarcity of controlled empirical work on the effects of novel predators on recipient communities and the mechanisms via which these effects play out (but see Lapiedra *et al.* 2018; Pringle *et al.* 2019). Such tests are needed, however, if we are to predict invasive species impacts, and improve conservation management (Sih *et al.* 2010a) and our understanding of how communities are structured via predator invasion (Sax *et al.* 2007).

Northern quolls (*Dasyurus hallucatus*) were, until recently, a common predator across northern Australia. They have declined over the last several decades, following the general decline in northern Australian mammals (Woinarski *et al.* 2015), thought to be driven by changes in grazing, fire, and predation regimes (Braithwaite & Griffiths 1994). More recently, the invasion of toxic invasive prey (cane toads, *Rhinella marina*) has resulted in dramatic, range-wide population declines in northern quolls (Shine 2010; Oakwood *et al.* 2016). Due to local extinction, northern quolls are now absent from large tracts of their former range and their ecological function as a medium-sized mammalian predator has been lost (Moore *et al.* 2019). For their conservation, northern quolls have recently been introduced to a number of offshore islands where they have never previously existed.

In 2017, a population of 54 northern quolls were introduced to a 25km^2^ island off the coast of north-western Northern Territory, Australia (Kelly 2019). Prior to this introduction, Indian Island (Kabarl) lacked mammalian predators, and large native reptilian predators had recently been reduced to near extinction by the invasion of cane toads. We take advantage of the introduction of northern quolls to a new island to directly test the effects of quolls as a novel predator on an island ecosystem and observe how native prey populations adjust to mitigate the impacts of their arrival. Since quolls are an ecologically novel predator on this island, we predict that this introduction may result in demographic effects (reduced survival and abundance) in invaded prey populations. If behavioural adjustments are able to reduce the demographic effects of a novel predator, we predict rapid behavioural changes in quoll-exposed melomys populations, such as changes in personality composition, foraging behaviour and responses to predator-scent, may manifest through time.

## Methods

### Introduction of northern quolls

In May 2017, 54 adult northern quolls were introduced to the north-eastern tip of Indian Island, Bynoe Harbor, Northern Territory, Australia (12°37’24.60”S, 130°30’0.72”E) to field test the conservation strategy of targeted gene flow (Kelly & Phillips 2016). Quolls are a voracious, opportunistic generalist predator (< 1.5k g; Oakwood 1997), and their introduction presented an opportunity to monitor the behavioural and demographic impacts on grassland melomys (*Melomys burtoni*), a native mammalian granivorous prey species (mean body mass 56 g, 5.6–103.7 g). Immediately prior to the introduction of quolls, we started monitoring populations of melomys in one woodland and two monsoon vine thicket plots in the vicinity of where quolls were to be released and radio tracked. After quolls were introduced and tracked it became immediately apparent that quolls were largely avoiding monsoon vine thicket sites and, since these sites would neither be effective “impact” or “control” sites, these sites were dropped from the on-going monitoring. Because these sites had to be dropped from our monitoring, we missed the opportunity to implement a robust Before-After Impact-Control design. For this reason, we only present data from before the introduction of quolls from one invaded site. Most of our data compare quoll-invaded (impact) versus quoll-free (control) sites over time, commencing within a few months of quoll arrival.

### Melomys population monitoring

To determine whether the arrival of a novel predator resulted in demographic impacts (population size and survival) to native prey species, we monitored four “impact”, quoll-invaded sites established in the north of Indian island in the vicinity of where quolls were released and three “control”, quoll-free sites established in the south of the island (Fig 1). Populations of melomys on Indian Island were monitored during four trips occurring immediately prior to the introduction of quolls in May (site 1) 2017, and after the introduction of quolls August 2017 (sites 2–7), April 2018 (sites 1–7), and May 2019 (sites 1–7).

**Figure 1.**
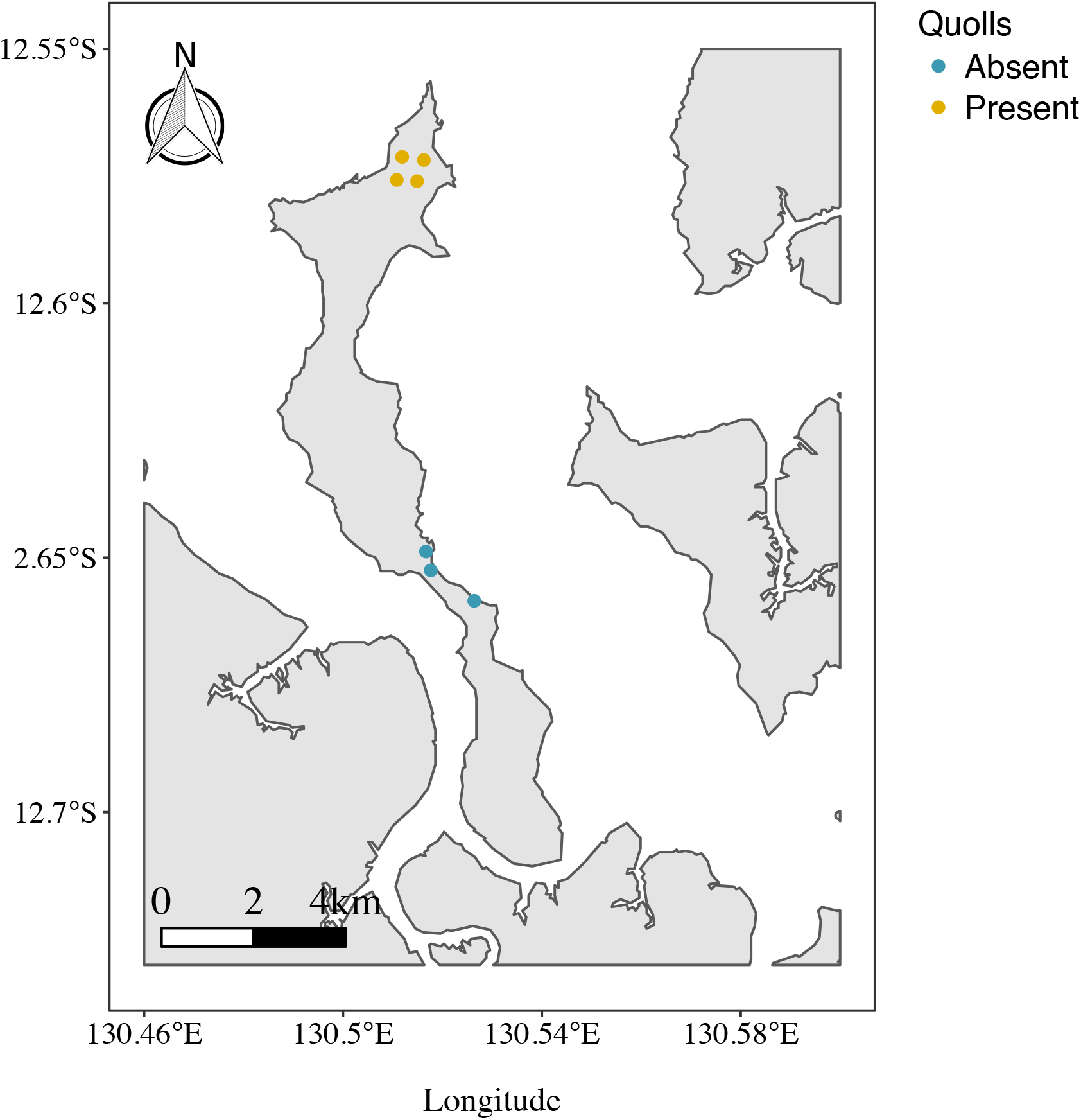
Map showing the arrangement of grassland melomys (*Melomys burtoni*) monitoring sites on Indian Island, Northern Territory, Australia. Quolls were present at the four monitoring sites in the north of the island and quolls were absent from the three monitoring sites in the south of the island for the duration of the study.

Melomys were monitored at seven independent 1ha (100 m x 100 m) plots (sites 1–7) spread out across Indian Island using a standard mark-recapture trapping regime designed for a monitoring project (Begg *et al.* 1983; Kemper *et al.* 1987). Sites in the north (quoll-invaded) and south (quoll-free) of the island were between 8.7 and 9.8km apart (Fig. 1; Table 1) and were composed of similar habitat types. The northern and southern sections of Indian Island are divided by mangrove habitat which is inundated at high tide. Cage and camera trapping as well as track surveys confirmed that quolls were present at the “impact” sites and absent from the “control” sites for the duration of the study (Jolly *et al.* unpub. data).

**Table 1.** Pairwise distance matrix between sites on Indian Island, Northern Territory, Australia. Quolls were present at sites 1–4 and quolls were absent at sites 5–7 for the duration of the study.

| Distance (m) | Site 1 | Site 2 | Site 3 | Site 4 | Site 5 | Site 6 | Site 7 |
| --- | --- | --- | --- | --- | --- | --- | --- |
| Site 1 |  |  |  |  |  |  |  |
| Site 2 | 270 |  |  |  |  |  |  |
| Site 3 | 260 | 350 |  |  |  |  |  |
| Site 4 | 400 | 300 | 250 |  |  |  |  |
| Site 5 | 8760 | 9030 | 8710 | 9000 |  |  |  |
| Site 6 | 8470 | 8730 | 8450 | 8070 | 300 |  |  |
| Site 7 | 9670 | 9920 | 9590 | 9820 | 1260 | 1500 |  |

Each of the seven monitoring sites consisted of 100 Elliott traps (Elliott Scientific Equipment, Upwey, Victoria) spaced at 10 m intervals in a 10 x 10 grid. Most trapping grids were open for four nights, however, the first trapping grid (site 1, May 2017) was open for six nights. After four trap nights, the majority of the melomys population had been captured at least once (Jolly *et al.* 2019). Traps were baited with balls of peanut butter, rolled oats and honey. These baits were replaced daily for the duration of each trapping session. Traps were checked for captures early each morning and all traps were cleared within two hours of sunrise.

Captured melomys were weighed (g) and sexed. Before release, each melomys was implanted with a microchip (Trovan Unique ID100). On successive mornings, all melomys were scanned (Trovan LID575 Handheld Reader), and any new individuals were microchipped. On the last morning of each trapping session, all melomys caught were retained for behavioural assays. Throughout the study 439 individual melomys were captured and given microchips (melomys caught per site: site 1 = 83; site 2 = 52; site 3 = 63; site 4 = 59; site 5 = 69; site 6 = 59; and site 7 = 54). Of these, 146 (33%) were caught on the final night of trapping and were retained for behavioural trials. Only large, healthy juveniles (*n* = 11), adult males (*n* = 58), and adult non-visibly pregnant females (*n* = 77) were retained for behavioural experiments. Melomys were retained in their respective Elliott traps and taken to the field station for diurnal husbandry. They were provided food and water *ad libitum* until 2 hours prior to testing. At this point, in an attempt to standardise hunger levels, access to food and water was removed. Indian Island is remote and uninhabited by humans, so all behavioural experiments were conducted in the field under near natural conditions (see Jolly *et al.* 2019 for detailed experimental procedures).

### Modified open field tests

We employed modified open field tests (also referred to as emergence tests: see Brown & Braithwaite 2004; López *et al.* 2005; Carter *et al.* 2013; Jolly *et al.* 2019) to assess boldness in grassland melomys and whether the arrival of a novel predator resulted in behavioural shifts in invaded populations. All open field tests were conducted on the night after the last trap night (night 5) and in opaque-walled experimental arenas (540mm x 340mm x 370mm). Experimental arenas were modified plastic boxes that had an inverted Elliott trap sized hole cut in one end and were illuminated by strings of red LED lights (Jolly *et al.* 2019). Each experimental arena had natural sand as substrate, and a rolled ball of universal bait (peanut butter, oats and honey) located both in the centre and along one wall of the arena (Jolly *et al.* 2019). After dark, Elliott traps containing a melomys were inserted into the hole in the side of each experimental arena and melomys were allowed to habituate for 10 min. At the start of each trial, Elliott trap doors were locked open—the inverted orientation of the trap prevented them from being triggered closed. Melomys were given 10 min to explore the open field arena. After 10 min, individuals were rounded back into their retreat (the Elliott trap) and a novel object (standard red, plastic disposable bowl) was placed at the end of the arena opposite the Elliott trap (Jolly *et al.* 2019). Melomys were then given a further 10 min to explore the arena and interact with the novel object. Elliott traps remained open during the open field tests and melomys could shelter and emerge from them under their own volition. All trials were recorded using a GoPro HERO 3. A previous study in this system determined that melomys showed repeatable behaviour between trials (boldness: *R* [± 95%CI] = 0.67 [0.47, 0.80], *P* < 0.001; emergence time: *R* [± 95%CI] = 0.73 [0.53, 0.83], *P* < 0.001; novel object: *R* [± 95%CI] = 0.61 [0.209, 0.974], *P* < 0.001; Jolly *et al.* 2019), therefore the data presented in this study were from a single behavioural trial of each animal (*n* = 146). Once trials were complete, each melomys was released at its point of capture.

To measure the boldness of individual melomys, we scored three behaviours typically associated with boldness and neophobia in rodents (Dielenberg & McGregor 2001; McGregor *et al.* 2002; Réale *et al.* 2007; Cremona *et al.* 2015): whether melomys fully emerged from their Elliott trap hide and entered the open arena during the 0–10 min period (scored 0 or 1, respectively); whether they fully emerged and entered the trial arena during the 10–20 min period (scored 0 or 1); and whether they interacted (touched) with the novel object that was placed in the arena during the 10–20 min period (scored 0 or 1). Videos were scored by a single observer who was blind to each melomys’ origin and identity. Because interacting with the novel object was predicated on a melomys’ willingness to emerge from their hide during the 10–20 min period, for analysis we combined their emergence during this period and interaction with the novel object into a single binary score: 0 (neophobic) = did not emerge or emerged but did not interact with novel object; or 1 (not neophobic): emerged and interacted with novel object.

### Seed removal plots

To assess whether the arrival of a novel predator affected the seed harvesting behaviour of granivorous melomys, we established seed removal plots at each site and sampled them each trapping session (night 6). After trapping and open field tests were conducted and melomys had been returned to their capture location, we set up 81 seed plots at each site by scraping away leaf litter with a shovel to create bare earth plots. These bare earth plots were created so that they were located in the centre between four Elliott traps within the 10×10 trapping grid. All seed plots were located randomly with respect to “distances to cover” but were all located on relatively open patches of ground. Sufficient within site replication (*n* = 81) significantly reduces the likelihood of distance to cover biasing population-level responses to seeds. Just before dark on the night of the seed removal experiment, we placed a single wheat seed in the centre of each bare earth plot. These seeds were either unscented, control seeds (*n* = 40) or predator-scented seeds that had been maintained in a sealed clip-lock bag filled with freshly collected northern quoll fur (*n* = 41). The placement of predator-scented and unscented seeds was alternated so that there was a chequered arrangement of scented and unscented seeds across the site. To ensure that the predator-scent was strong enough to be detected by melomys, along with the predator-scented seeds, we also placed a few strands of quoll fur around the predator-scented seeds. Before light the next morning, we returned back to each plot and counted the number of seeds of each scent-type that were removed from the plot. Melomys are the only nocturnal granivorous animal that occurs on Indian Island, and to avoid diurnal granivorous birds from removing seeds we conducted this experiment during the night only.

### Wildfire on northern Indian Island

Immediately following our monitoring and experiments in August 2017, a wildlife broke out on northern Indian Island in the vicinity of the four quoll-invaded sites and burnt through all of the sites. Because of this, our experimental design is confounded by the fact that all of our quoll-invaded sites were burnt, and all of our quoll-free sites were unburnt. Fire is a regular disturbance in this landscape (Andersen *et al.* 2005), and previous work has shown little effect of fire on abundance, survival or recruitment of grassland melomys (Griffiths & Brook 2015; Liedloff *et al.* 2018). Nonetheless, this confound exists and we proceed with caution when interpreting the effects of quolls on population size and survival of melomys.

### Statistical analysis

During trapping sessions we identified individual melomys that were captured at each site by their unique microchips. Because melomys on Indian Island have very small home ranges (tending to be caught in the same or adjacent traps throughout the trapping period: Jolly *et al.* unpub. data) and since we never observed captures of melomys marked at other sites (Jolly *et al.* unpub. data), we treated each site as independent with regard to demographics and behaviour (Table 1).

To estimate between-session survival, we analysed the mark-recapture data to estimate recapture and survival rates using Cormack-Jolly-Seber models in program MARK. At each site, there were three primary trapping sessions of four nights, for a total of 12 time intervals in the input file. Because quolls prey on melomys, we hypothesised that survival rates of melomys would be lower between trapping sessions at sites with quolls than at sites without quolls. We included two groups, quoll-free (control) and quoll-invaded (impact), in the input file. We ran a series of models in MARK to test the following *a priori* hypotheses: (1) survival rates between sessions are lower at quoll-free sites than at quoll-invaded sites; (2) survival rates are lower between sessions than within sessions, but are unaffected by quolls; (3) survival is constant through time; and (4) survival varies through time. All candidate models were ranked according to their AICc values and associated AIC weights (Burnham & Anderson 1998). Models with AICc values < 2 were considered to be well supported by the data (Burnham & Anderson 1998). We used Akaike’s Weights, which are proportional to the normalized, relative likelihood of each model, and to determine which of these models was most plausible (Buckland *et al.* 1997).

To test whether the presence of quolls impacted melomys population size, we used a hierarchical model in which population size was made a function of quoll presence/absence, capture session, and the interaction between these factors. Population size at each site during each session is estimated in this process, and we fitted this model in a Bayesian framework. Our observations consisted of a capture history for each observed individual over the number of nights at each site for each trapping session. We denoted the number of individuals at site *s* during session *k* as *N_ks_*. To estimate *N_ks_* we used a closed population mark-recapture analysis in which each individual, *i*, was either observed, or not (*O_iks_*), according to a Bernoulli distribution:

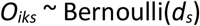

Where *d_s_* denotes the expected detection probability within session *s*. Our previous MARK analysis found clear evidence for variation in detection probability across sessions, but detection probabilities of melomys on Indian Island had previously been found not to vary measurably between individuals nor to change over time within a trapping session (Jolly *et al.* 2019). Thus, we made detection probability a function of session according to:

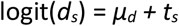

Where *μ_d_* is the expected detection probability in the first session, and *t_s_* denotes the (categorical) effect of session on detection.

We used the “data augmentation” method (Tanner & Wong 1987; Royle *et al.* 2007; Kery & Schaub 2011) in combination with this detection probability to estimate *N_ks_* for each site per session (site.session). Using this approach, the data were ‘padded’ to a given size by adding an arbitrary number of zero-only encounter histories of ‘potential’ unobserved individuals. The augmented dataset was then modelled as a zero-inflated model (Royle *et al.* 2007) which changes the problem from estimating a count, to estimating a proportion. This was executed by adding a latent binary indicator variable, *R_iks_*, (taking values of either 0 or 1) to classify each row in the augmented data matrix as a ‘real’ individual or not, where *R_iks_*,~ Bernoulli(*Ω_ks_*). The parameter *Ω_ks_* is the proportion of the padded population that is real, and *N_ks_* = Ω*_i_ R_iks_*.

We then made *Ω_ks_* (which scales with population size) a function of quoll presence/absence, *q_c_*; session, *b_k_*; and the interaction between the two:

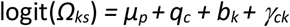

The model was fitted using Bayesian Markov Chain Monte Carlo (MCMC) methods and minimally informative priors (Table 2) within the package JAGS (Plummer *et al.* 2017) using R (R Core Team 2019). Parameter estimates were based on 30,000 iterations with a thinning interval of 5 following a 10,000 sample burn-in. Three MCMC chains were run, and model convergence assessed by eye, and using the Gelman-Rubin diagnostic (Gelman & Rubin 1992a, 1992b).

**Table 2.** Results of Cormack-Jolly-Seber analyses used to compare survival (*Phi*) and recapture (*p*) probabilities of grassland melomys (*Melomys burtoni*) on Indian Island, Northern Territory, Australia. The symbols ‘.’ and ‘*t*’ refer to constant and time, respectively, while ‘g’ denotes the two groups (quoll free versus quolls present). Table shows AIC values and associated AIC weights, model likelihood, number of parameters (*N*), and model deviance. The term ‘w/b’ indicates that within trapping session survival rates (s_1_-s_3_, s_5_-s_7_, s_9_-s_11_) were constant and equivalent, and different to the between trapping session survival rates (s_4_, s_8_). The term ‘group w/b’ is as above, except that between trapping session survival rates differed between the two groups.

| Model | AICc | Delta AICc | AICc Weights | Model Likelihood | N | Deviance |
| --- | --- | --- | --- | --- | --- | --- |
| Phi (group w/b) p(t) | 1688.629 | 0 | 0.92162 | 1 | 16 | 477.5224 |
| Phi (w/b) p(t) | 1693.562 | 4.933 | 0.07823 | 0.0849 | 14 | 486.6202 |
| Phi (group w/b) p(g x t) | 1701.116 | 12.4873 | 0.00179 | 0.0019 | 27 | 466.705 |
| Phi (group w/b) p(g x t) | 1703.132 | 14.503 | 0.00065 | 0.0007 | 7 | 510.5963 |
| Phi (t) p (g*t) | 1718.863 | 30.2344 | 0 | 0 | 32 | 473.631 |
| Phi (t) p (.) | 1719.862 | 31.2339 | 0 | 0 | 12 | 517.0641 |
| Phi (t) p (g) | 1720.751 | 32.1229 | 0 | 0 | 13 | 515.8843 |

To assess whether the introduction of quolls affected the behaviour of melomys populations, we divided the responses of melomys in open field tests into two independent response variables: whether individuals emerged or not during the 0-10 min period (binomial: 0 or 1); and whether individuals emerged and interacted with the novel object or not during the 10-20 min period (binomial: 0 or 1). We used generalised linear mixed-effects models with binomial errors and a logit link to test the effect of quoll presence (two levels: quolls present and quolls absent) and trapping session (continuous), with site included as a random effect, on the behavioural response variables. *P*-values were obtained by likelihood ratio tests of the full model with the effect in question against the model without the effect. This analysis was performed using R with the *lme4* software package (R Core Team 2019).

To assess whether the numerical impact of quolls on melomys affected the seed harvesting rate of invaded melomys populations, we first examined the relationship between melomys population size (estimated above) and the total number of control (unscented) seeds harvested from each site. Here we used a simple linear model with number of seeds harvested as a linear function of population size, quoll presence/absence and the interaction between these effects. To test whether there was an additional effect of quoll presence, beyond their effect on population size, we defined a new variable, Δ_ks_, as the difference in seed take between scented and unscented treatments within each site.session. Here any effect of melomys density is cancelled out (because density is common to both treatments within each site.session). Thus, we fitted a model in which Δ_ks_ is a function of quoll presence/absence, session and the interaction between these effects. These analyses were performed using R version 3.3.2 (R Core Team 2019).

## Results

### Effect of novel predator on survival

When we assessed the impact of quolls on melomys survival between trapping sessions the best supported model was one in which survival rates between sessions were lower at quoll-invaded sites than at quoll-free sites, and recapture rates were session-dependent (Table 2). All other models were more than 4 AIC units from this best model, and so clearly inferior descriptions of the data. From the best-supported model, estimates of apparent survival (*S*) for the intervals between the capture sessions were substantially higher at quoll-free sites (*S_2017–2018_* = 0.368; *S_2018–2019_* = 0.225) than at quoll-invaded sites (*S_2017–2018_* = 0.207; *S_2018–2019_* = 0.091; Fig. 2). The differing survival probability between sessions is largely explained by the time difference between intervals (2017–2018 = 9 months vs. 2018–2019 = 13 months; Fig. 3).

**Figure 2.**
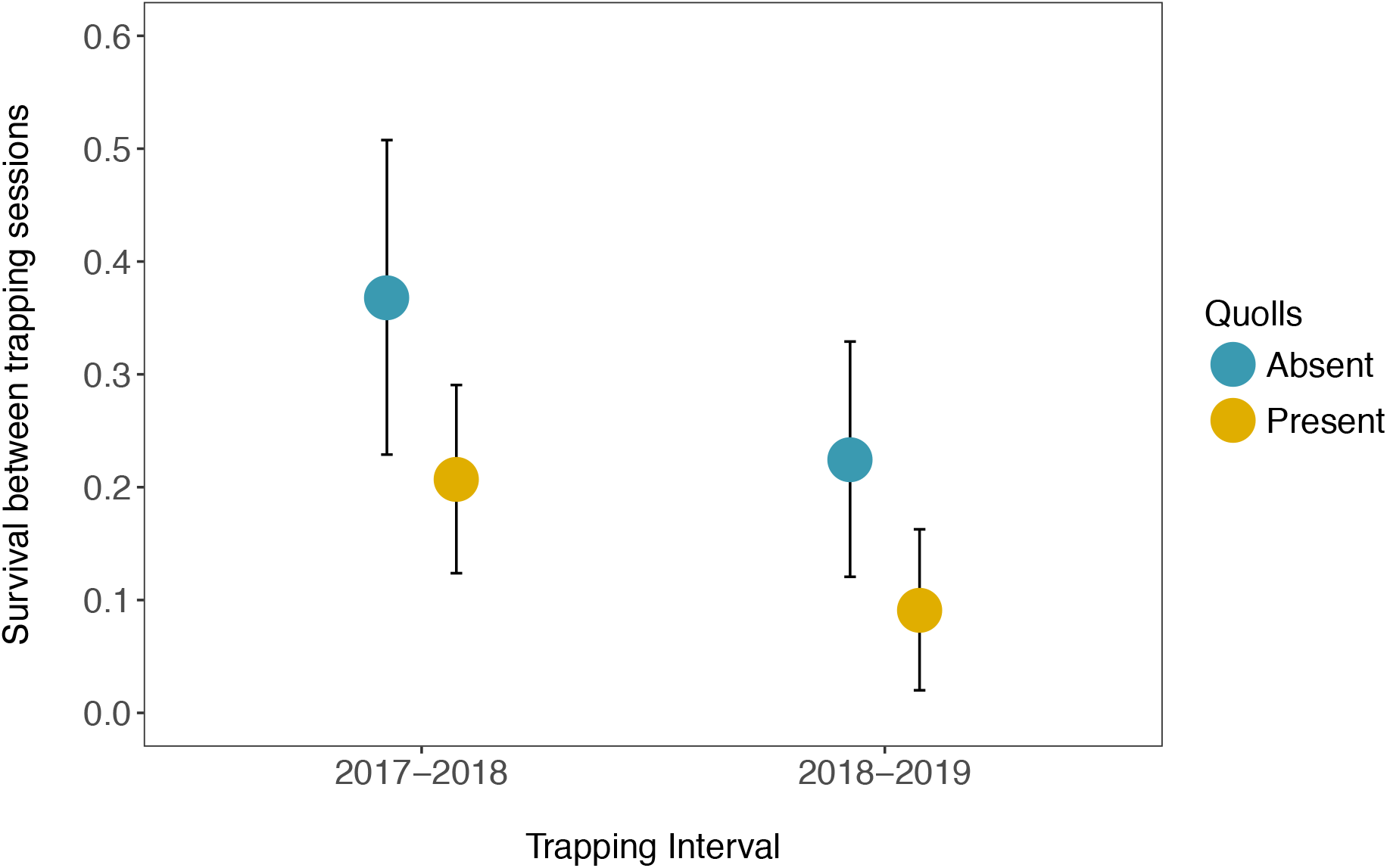
Between trapping session survival (± 95% CI) of grassland melomys (*Melomys burtoni*) on Indian Island in quoll-invaded (*n* = 4) and quoll-free (*n* = 3) populations on Indian Island, Northern Territory, Australia.

**Figure 3.**
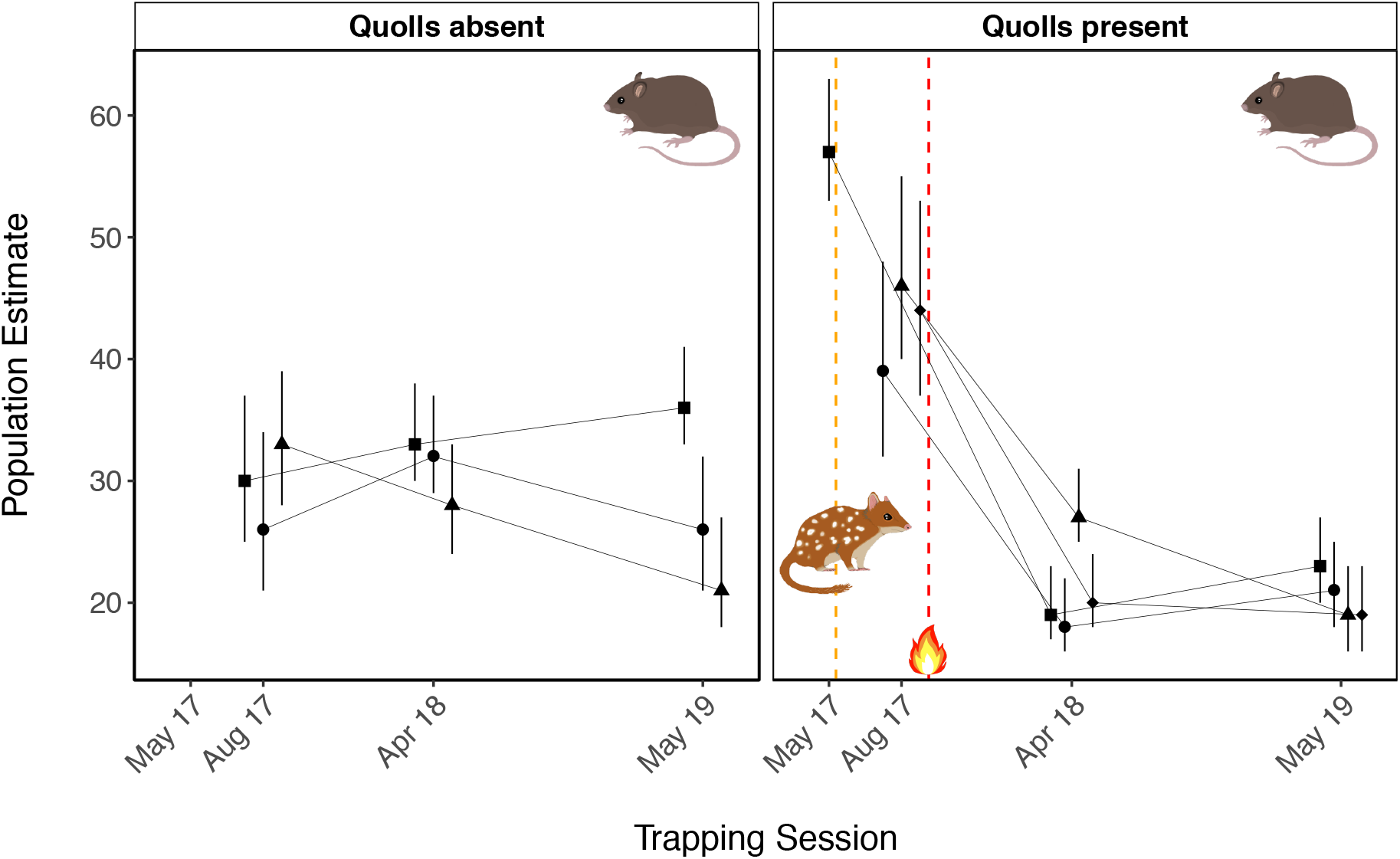
Posterior mean population sizes (*N_ks_* ± 95% CI) for quoll-invaded and quoll-free populations of grassland melomys (*Melomys burtoni*) on Indian Island, Northern Territory, Australia. The orange dotted vertical line denotes the timing of the introduction of quolls. The red dotted vertical line denotes the timing of an unplanned fire that burnt through the quoll-invaded sites. In each predator treatment, different sites are denoted by different shaped points. Estimates assume closure of the population within each session and detection probability that varies across sessions.

### Effect of novel predator on population size

Populations of melomys declined dramatically in quoll-invaded sites in the year following their introduction but not in quoll-free sites (Fig. 3). We observe a strong negative interaction between the presence of quolls and trapping session in 2018 (mean = −1.194, 95% credible interval [−1.732, −0.665]) and 2019 (mean = −1.097, 95% confidence interval [−1.652, −0.551]; Fig. 3; Table 3).

### Effects of novel predator on prey behaviour

For the proportion of melomys emerging in open field tests during the 0–10 min period, there was a significant interaction between quoll presence and trapping session (χ^2^ (5) = 4.386, *P* = 0.04; Fig. 4). There was no interaction between quoll presence and trapping session for the proportion of melomys emerging and interacting with the novel object during 10–20 min period (χ^2^ (5) = 2.567, *P* = 0.109; Fig. 4). The model without this interaction, however, revealed a significant effect of quoll presence, with fewer melomys emerging from hiding and interacting with the novel object during the 10–20 min period of open field tests from sites where quolls were present than from sites where quolls were absent (χ^2^ (5) = −4.696, *P* < 0.001; Fig. 4).

**Table 3.** Model parameters and their priors including prior distributions, standard deviation, estimated posterior means and their 95% credible intervals. N denotes normal probability distribution with mean and standard deviation.

| Model Parameters |  |  |  |  |
| --- | --- | --- | --- | --- |
| Name for parameter | Parameter | Prior (mean, SD) | Posterior mean | 95% credible intervals |
| **Detection:** |  |  |  |  |
| Intercept for detection | $\mu_d$ | N (0, 2.71) | -0.94 | -1.12, -0.76 |
| Effect of session 2 on detection | $t_2$ | N (0, 2.71) | 0.59 | 0.33, 0.85 |
| Effect of session 3 on detection | $t_3$ | N (0, 2.71) | 0.46 | 0.18, 0.73 |
| **Population size:** |  |  |  |  |
| Intercept for Omega | $\mu_p$ | N (0, 2.71) | -0.91 | -1.22, -0.58 |
| Quoll Presence | $r_2$ | N (0, 2.71) | 0.70 | 0.31, 1.09 |
| Trapping Session 2 | $b_2$ | N (0, 2.71) | 0.06 | -0.36, 0.48 |
| Trapping Session 3 | $b_3$ | N (0, 2.71) | -0.10 | -0.53, 0.34 |
| Interaction 1 [Quoll Presence * Trapping Session 2] | $\gamma_{2,2}$ | N (0, 2.71) | -1.19 | -1.73, -0.67 |
| Interaction 2 [Quoll Presence * Trapping Session 3] | $\gamma_{2,3}$ | N (0, 2.71) | -1.10 | -1.65, -0.55 |

**Figure 4.**
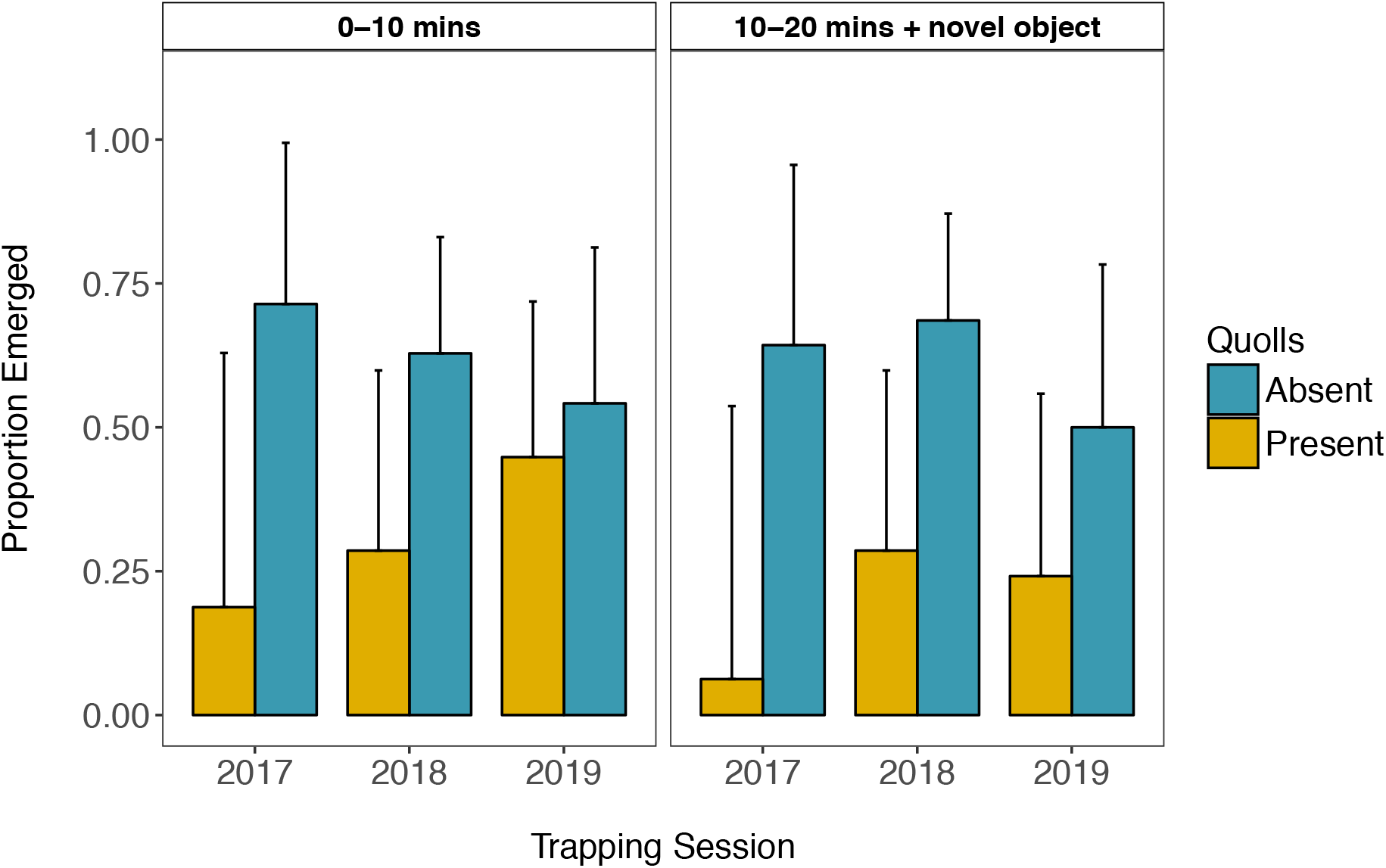
Mean proportion (± 95% CI) of grassland melomys (*Melomys burtoni*) emerging from hiding during open field tests from quoll-invaded sites in 2017 (*n* = 16), 2018 (*n* = 28) and 2019 (*n* = 29), and quoll-free sites in 2017 (*n* = 14), 2018 (*n* = 35) and 2019 (*n* = 24) on Indian Island, Northern Territory, Australia.

### Effects of novel predator on seed harvesting and predator-scent aversion

Although there was no interaction between melomys density and quoll presence (*t*_18_ = −0.251, *P* = 0.805; Fig. 5), there was a very clear positive relationship between melomys density and seed take (*t*_18_ = 5.112, *P* < 0.001; Fig. 5) and a clear negative relationship between quoll presence and seed take (*t*_18_ = −2.344, *P* = 0.031; Fig. 5).

**Figure 5.**
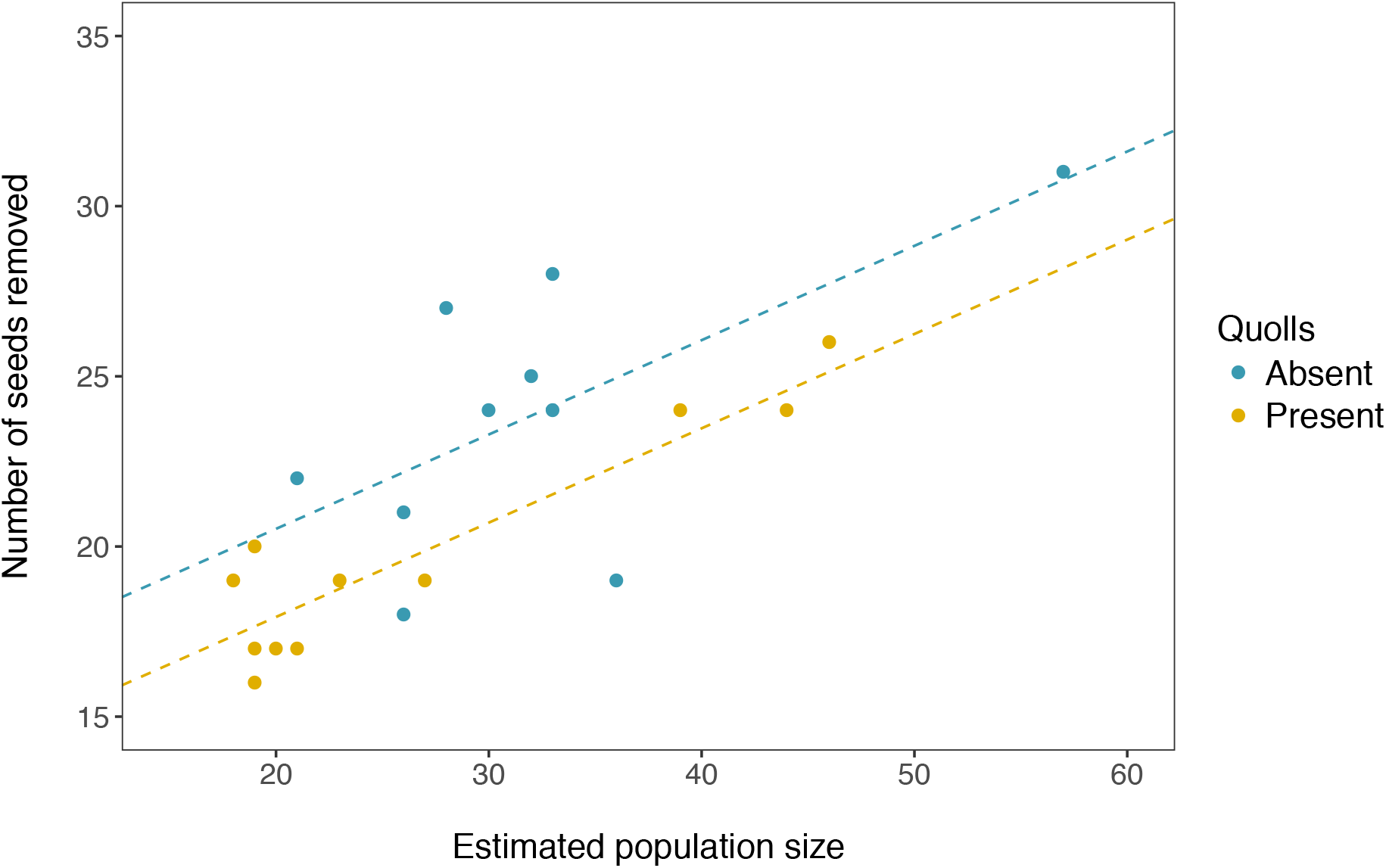
Effect of estimated population size on the number of control, unscented seeds removed from seed plots (*n* = 21) in quoll-invaded and quolls-free sites. Dotted lines denote the effect of quoll presence on seed removal rate.

When we looked at the difference in seed take (Δ_ks_) between scent treatments within site.session, a striking pattern emerges, in which there is a clear interaction between the presence of quolls and session (*F*_3,17_ = 18.61, *P* < 0.001; Fig. 6).

**Figure 6.**
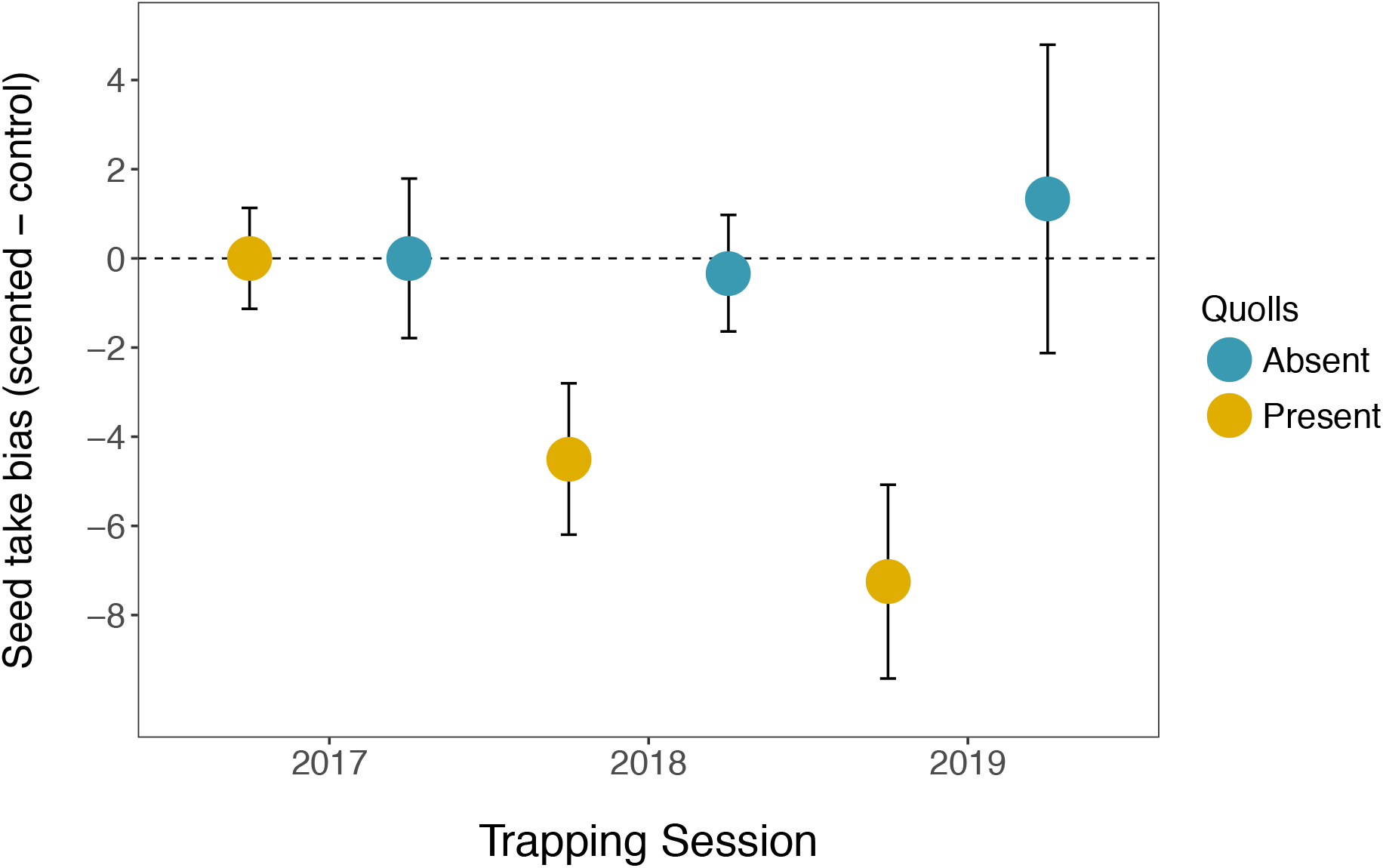
Mean (± 95% CI) difference (Δ) between the number of predator-scented seeds and control, unscented seeds removed by melomys from quoll-invaded (*n* = 3; 2017 & *n* = 4; 2018-19) and quoll-free (*n* = 4; 2017 & *n* = 3; 2018-19) sites during each trapping session.

## Discussion

The introduction of northern quolls to Indian Island was associated with lowered survival and an apparent drop in population size in quoll-invaded melomys populations. This numerical effect on melomys density had an impact on seed predation rates, because seed take is strongly associated with the density of melomys in this system. This is a classic trophic cascade: predation suppresses herbivore density, which reduces the pressure that herbivores place on primary producers. Our study, however, also reveals an additional, subtler, cascade effect; driven by altered prey behaviour rather than by altered prey density.

Within months of quolls appearing on the island, invaded populations of melomys were significantly shyer than nearby, predator-free populations of conspecifics. This rapid but generalised response to a novel threat appears to have had a subtle effect on seed predation rates: when we examine unscented seeds, per capita seed take is slightly lower in quoll-invaded populations. This generalised response appears to have been supplemented over time with more threat-specific antipredator behaviours. Although the willingness of predator-exposed melomys to emerge from shelter (i.e. boldness) converged through time with that of predator-free melomys, predator-exposed melomys continued to be more neophobic than their predator-free conspecifics throughout the study. Meanwhile, predator-scent aversion, as evidenced by seed plots, steadily increased over time. Presumably the significant and novel predation pressure induced by the introduction of quolls resulted in selection on behaviour and/or learning in impacted rodent populations, allowing them to fine-tune their behavioural response (decrease general shyness, but maintain neophobia, and respond to specific cues) as the nature of the threat became clearer. These changing behavioural responses imply a generalised reduction in seed take that also becomes fine-tuned over time, with high risk sites (those that smell of predators) ultimately displaying substantially lower seed take than low risk sites. Thus, we see a reduction in seed take resulting in a fine-scaled aversive response varying on a spatial scale measured in the tens of metres.

Although our study documented dramatic population declines in predator-invaded melomys populations, and we are assigning the causation of these declines to the introduction of quolls, we need to address the confounding factors that may affect how we interpret our results. Firstly, there is an inherent and unavoidable spatial confound in our study system driven by the location of our study sites. We cannot exclude the possibility that some of the population change we observe in our predator-invaded populations could also be due to the population naturally declining towards sustainable levels unrelated to the addition of a novel predator. It is possible that, by chance, when we started monitoring populations of melomys, populations at northern sites were at a population peak and were naturally cycling towards sustainable levels, while southern populations were stable. However, although we cannot rule this out, such between population differences would be expected to be driven by differences in resource availability between the locations (e.g. Dickman *et al.* 1999; Russell & Ruffino 2012). We believe this is unlikely in our study system, given the relatively close proximity of our sites (<10 km) and the spatially homogenous climatic conditions that govern the wet-dry monsoonal tropics of northern Australia. Rodent population cycles in the Australian wet-dry tropics appear to be primarily driven by annual differences in rainfall between wet seasons, rather than spatial differences within years (Madsen & Shine 1999). For this reason, we suspect natural population cycles are unlikely to explain the population change differences we observe in this study.

Additionally, there is the unplanned, confounding factor of the fire that burnt through northern Indian Island after completion of our population monitoring in 2017. Such fires are commonplace in the Australian wet-dry tropics (Russell-Smith & Yates 2007); a regular disturbance that is often rapidly offset by the annual monsoon driven wet season. Since our sites are composed of grass-free woodland, the fire that burnt through them mostly burnt leaf-litter (though it reached the mid-storey in other parts of the island). While this likely reduced the short-term availability of food and cover for melomys, it is unlikely to directly explain the demographic effects we observed. A previous study investigating the effect of fire regimes on native mammals in savanna woodland in Kakadu National Park, Northern Territory was unable to detect an effect of fire frequency or intensity on the survival or recruitment of grassland melomys, despite finding fire impacts in all other co-occurring native mammals studied (Griffiths & Brook 2015). Interestingly, even in a system where fire is much more infrequent and significantly more intense (e.g. mesic habitats of eastern Australia), grassland melomys were found to be relatively unaffected by a wildfire that caused significant impacts to a co-occurring native rodent, and any demographic impacts felt by melomys were entirely absent within months of the fire (Liedloff *et al.* 2018). Additionally, the most dramatic behavioural difference (boldness and neophobia) between quoll-invaded and quoll-free sites was observed immediately prior to the occurrence of the fire (early August *vs.* mid-August 2017). For the behavioural changes we observed that were potentially confounded by fire, such as predator-scent aversion, we would expect to see these effects decreasing with time since fire if fire was driving this response, instead we see the opposite trend. Finally, if food had become strongly limiting as a consequence of the fire, we would expect to have observed an increase in seed take in the burned (quoll-invaded) sites, instead we saw a decrease. For these reasons, we suspect the fire was unlikely to be directly responsible for the demographic effects to melomys we observed, and fire cannot in any way explain the response we observed to quoll-scented seeds. We, therefore, believe our interpretation of these changes as being driven mostly by the addition of a novel predator to the system is the most parsimonious and globally coherent interpretation of the data.

Predation is a pervasive selective force in most natural systems, driving evolutionary change in prey morphology, physiology, life history and behaviour. Unlike morphology and physiology, however, the labile nature of behaviour makes it a particularly powerful trait for rapid response in a changing world (Réale *et al.* 2007; Sih *et al.* 2010b; Dall & Griffith 2014). Behavioural comparisons of wild populations exposed to differing predation regimes provides some support for the prediction that reduced boldness would be selected for under high predation scenarios (Åbjörnsson *et al.* 2004; Bell 2005; Brydges *et al.* 2008) and that the appearance of novel predators can result in bold individuals becoming shyer (Niemelä *et al.* 2012), however, the opposite pattern of response can also occur (Brown *et al.* 2005; Urban 2007) or behavioural phenotypes can be unrelated to predation regime (Laurila 2000; Carlson & Langkilde 2014). Interestingly, a number of studies have demonstrated that individuals from high-predation areas were quicker to emerge (Harris *et al.* 2010) and were bolder and more aggressive (Bell & Sih 2007; Dingemanse *et al.* 2007) than predator-naïve conspecifics. Although we found the opposite pattern to this immediately following the arrival of a novel predator, by the second year after predator introduction we found the boldness of melomys converging with that of predator-free populations. Thus, it is clear that the behavioural composition of these populations are dynamic, and it seems likely this dynamism (and perhaps the capacity of the prey species to identify specific threats) may explain some of the variation between earlier studies.

Although boldness may change over time, neophobia, as a generalised adaptive response to predation pressure, is now well supported across a number of studies (Crane *et al.* 2019). Individuals living under high predation risk scenarios have been shown to typically display generalized neophobia (Brown *et al.* 2015; Elvidge *et al.* 2016), and neophobia can increase the survival of predator-naïve individuals in initial encounters with predators (Ferrari *et al.* 2015; Crane *et al.* 2018). Certainly, in our study, predator-exposed melomys were significantly more neophobic than their predator-free conspecifics; an effect maintained throughout the study. Despite reduced survival, significant population declines, and clear behavioural changes in invaded populations, it is impossible to determine with certainty from our data whether changes in the behaviour of predator-invaded melomys populations are the result phenotypic plasticity (learning) or natural selection. The low between trapping session survival of melomys in quoll-invaded populations means few individuals survive between sessions, so natural selection is a possibility, and selection on these behavioural traits is potentially very strong. Although behavioural changes in predator-invaded populations have been documented in a few systems where predator introductions have been staged and experimentally controlled (Lapiedra *et al.* 2018; Blumstein *et al.* 2019; Cunningham *et al.* 2019; Pringle *et al.* 2019), elucidating whether these observed changes arise because of behavioural plasticity or natural selection can be exceptionally difficult. Rapid behavioural responses of vulnerable prey to recovered predators has been observed in a single prey generation, presumably due to behavioural plasticity (Berger *et al.* 2001; Cunningham *et al.* 2019). Similarly, behavioural adjustments to an introduced predator have been observed as a result of natural selection on advantageous behavioural traits (Lapiedra *et al.* 2018). In this study, although we had measures of individual behaviour, our between session recapture rates of these individuals was sufficiently low that we had no longitudinal data on the behaviour of individuals to test whether individuals were altering their behaviour or whether natural selection was resulting in population-level change. It thus remains possible (and quite likely) that both mechanisms were in play.

Although northern quolls represent a novel predator to melomys on Indian Island, the two species’ shared evolutionary history on the northern Australian mainland may provide some explanation as to why this staged introduction resulted in rapid, finely-tuned behavioural adjustment in melomys, rather than extinction. Isolation from predators can rapidly result in the loss of antipredator behaviours from a prey species’ behavioural repertoire (Blumstein & Daniel 2005; Jolly *et al.* 2018a), dramatically increasing an individual’s susceptibility to predation following the introduction of either predator or prey (Carthey & Banks 2014; Jolly *et al.* 2018b). But such outcomes are not inevitable: length of isolation, co-evolutionary history, degree of predator novelty, density-dependent effects, population size, and pre-existing predator-prey associations (Berger *et al.* 2001; Blumstein 2006; Banks & Dickman 2007; Sih *et al.* 2010a; Carthey & Banks 2014) are all likely to be hugely influential in determining whether an invaded population adjusts to the invader or proceeds towards extinction. Recently, a conservation introduction of Tasmanian devils to an island previously lacking them found that their possum prey rapidly adjusted their foraging behaviour to accommodate this newly arrived predator (Cunningham *et al.* 2019). Despite possums having lived on the island in isolation from devils since the 1950s, presumably, their long evolutionary history together on mainland Tasmania had them primed to respond to this predatory archetype (Sih *et al.* 2010a; Carthey & Banks 2014; Cunningham *et al.* 2019). This shared evolutionary history is likely responsible for both possums’ and melomys’ ability to rapidly mount appropriate antipredator responses to the introduction of these predators. The predators are novel within an individual’s lifetime, but the individual’s ancestors have encountered them before.

Although our results suggest that invaded melomys populations are beginning to adjust to the presence of northern quolls as a novel predator on Indian Island, there has been no sign of demographic recovery from the addition of this predation pressure on the island. Data from our seed removal experiment clearly demonstrated that the function of melomys as seed harvesters and dispersers scales with density. Trophic cascades resulting from the addition and loss of predators from ecosystems has been observed in a number of systems globally (Ripple *et al.* 2001; Terborgh *et al.* 2001; Estes *et al.* 2011), and the results can profoundly shape entire systems. As the only rodent and the dominant granivore in this system, while melomys populations may or may not go extinct as a result of quoll invasion, their reduced abundance and weakened ability to harvest and disperse seeds may have yet to be observed, longer-term consequences for the vegetation structure and ecosystem function of Indian Island (McConkey & O’Farrill 2016). Currently, grass is a rare vegetation feature on Indian island (though it is a dominant feature of savanna woodlands generally), and this is quite possibly a result of the high density of melomys on this (previously) predator-free island. The presence of quolls may well change that, as both numerical and behaviour responses of melomys cascade down to the grass community.

Empirical research on the effects of novel predators on recipient communities under controlled conditions on a landscape-scale is exceptionally difficult and remains relatively rare. The introduction of threatened predators to landscapes from which they have been lost (Cunningham *et al.* 2019) or where they are entirely novel (Lapiedra *et al.* 2018), however, provides a unique opportunity to observe how naïve prey can respond to novel predators, and the mechanisms by which predators can structure communities. Our study provides empirical support that some impacted prey populations can adjust rapidly to the arrival of a novel predator via a generalised behavioural response (decreased boldness) followed by development of a species-specific antipredator response (behavioural fine-tuning). The arrival of the novel predator appears to have set off a trophic cascade that was likely driven, not only by changed prey density, but also by changed prey behaviour. Thus, rapid adaptive shift may allow prey populations to persist, but large-scale, system-wide changes may still follow.

## Data accessibility

Data are available online: https://doi.org/10.5281/zenodo.3936465

## Supplementary material

Scripts and code are available online: https://doi.org/10.5281/zenodo.3936465

## Acknowledgements

Thanks to Kenbi Traditional Owners (Raylene and Zoe Singh) for land access permission and Kenbi Rangers for assistance in the field. Special thanks to Kenbi Rangers Brett Bigfoot, Rex Edmunds, Jack Gardner, Ian McFarlane, Dale Singh, and Rex Sing for continued field assistance throughout this project. Thanks to Kenbi Ranger Co-ordinator Steven Brown for logistical support in the field. Thanks to Alana de Laive for graphic design of figures. Thanks to Ella Kelly and Naomi Indigo for logistical and moral support on the island. In kind support was provided by Kenbi Rangers and the Northern Territory Government Department of Environment and Natural Resources, Flora and Fauna Division (via GRG). CJJ was supported by an Australian Postgraduate Award, the Holsworth Wildlife Research Endowment and David Hay Postgraduate Writing Up Award. We thank two anonymous reviewers and Professor Réale for their comments via Peer Community in Ecology, which greatly improved the manuscript. Version 6 of this preprint has been peer-reviewed and recommended by *Peer Community In Ecology* (https://doi.org/10.24072/pci.ecology.100057).

